# The Effect of 1-MCP on grain size and yield-related traits of *Eragrostis tef*

**DOI:** 10.1101/2023.03.21.533369

**Authors:** Fano Dargo Girmay, Richard J. Flavel, Heather M. Nonhebel

**Affiliations:** School of Science and Technology, University of New England, Armidale, NSW, 2351, Australia; Dept Agronomy and Soil Science, Faculty Science, Agriculture, Business and Law, University of New England, Armidale, NSW, 2351, Australia

**Keywords:** grain size, TGW, 1-MCP, teff, panicle, yield-related traits

## Abstract

Teff is an Ethiopean cereal that has the potential for increased global production. However, as a very small-seeded grain, it experiences high levels of post-harvest loss. As ethylene is reported to have a negative effect on grain size, this study sought to test whether 1-methylcyclopropene (1-MCP), an ethylene action inhibitor, is a suitable molecule to improve grain size of teff as well as to determine the optimum application time and concentration. The 1-MCP treatment was evaluated on well-watered plants as well as plants subjected to moderate water stress, with the stress treatment imposed one week before the expected flowering date. Application of 1 PPM 1-MCP to individual panicles improved grain size and thousand-grain weight (TGW) by 22% and 29% under moderate water stress and 10% and 16% under well-watered conditions, respectively. It also increased yield per panicle. However, when applied to whole plants it was not effective, due to the extended flowering time of teff and differing responses of panicles at different developmental stages. These observations suggest that genetic modification leading to lower production of endogenous ethylene could benefit grain size in teff. However, 1-MCP treatment is unlikely to be effective in the field.

## Introduction

Teff [*Eragrostis tef (Zucc.*) Trotter] originated and was domesticated in Ethiopia between 4000-1000 BC (Vavilov, 1951). In Ethiopia, teff is a staple food crop and is used in the form of “injera” (sourdough flatbread) (Gebru *et al*., 2019). Teff has several health benefits, including the absence of gluten, low glycemic index, high mineral content (Fe, Ca, Cu, Zn, P, and K), high protein and fibre content, as well as a well-balanced composition of essential amino acids (Baye, 2014; Dereje *et al*., 2019; Spaenij-Dekking *et al*., 2005). As a result of its health significance as well as climate resilience and resistance to biotic and abiotic stress, the production of teff for human consumption is now expanding outside of Ethiopia (Cheng *et al*., 2017; Crymes, 2015; Ketema, 1997). However, the grain size of teff is very small and because of this the post-harvest loss is very high, estimated at 40.9% (22.8% at the farmer level, 2.2% at the Rural assemblers/Zone traders, 8.5% at the Wholesalers, and 7.4% at the Retailers/Millers) (FAO, 2018). The small grain size also results in difficulty in seed-handling at the time of planting, and seed contamination with soil, sand, weed, grass, and straw at the time of threshing and harvesting (Chanyalew *et al*., 2013; Fufa *et al*., 2013; Van Delden *et al*., 2010). Thus, increasing grain size is vital to minimize post-harvest loss and other production constraints within Ethiopia as well as increasing the attractiveness of the crop for international production.

In their review, Kesavan *et al*. (2013) have described the importance of phytohormones in determining seed size and as a target for breeding. Endogenous cytokinin (CK), indoleacetic acid (IAA), and abscisic acid (ABA) are positively correlated with grain-filling rate and duration (Li *et al*., 2020; Wang *et al*., 2006; Wei *et al*., 2019; Yang *et al*., 2003; Yang *et al*., 2004; Zhang *et al*., 2016). The application of CK, auxin, and ABA during the early grain-filling stage also has a positive effect on grain size and yield components. Application of CK to rice, wheat, maize, and triticale increased the grain weight, grain-filling period, and rate (Basunia and Nonhebel, 2019; Li *et al*., 2020; Panda *et al*., 2018; Sivakumar *et al*., 2001). Similarly, IAA treatment of rice enhanced the grain filling rate and grain weight (Yang *et al*., 2003; Zhang WY *et al*., 2016). Foliar spray of rice and wheat with ABA also improved grain filling rate and thousand-grain weight (TGW) (Yang *et al*., 2006; Yang *et al*., 2004). The effects of altered expression or mutations of hormone-related genes also confirm the importance of phytohormones for grain size.

In contrast to the positive effects of CK, IAA, and ABA on grain size, the high production of ethylene has frequently been linked to reduced grain filling rate, grain filling period, and grain weight, as well as increased kernel abortion, ultimately resulting in reduced grain size and yield (Hussain *et al*., 2019; Wang *et al*., 2012; Yang *et al*., 2004; Yang *et al*., 2014). Several types of abiotic stress (salt, heat, and water-deficit) promote ethylene production (Hays *et al*., 2007; Hussain *et al*., 2019; Yang *et al*., 2004; Yang *et al*., 2014), thus mediating or exacerbating the negative effect of stress on grain size. The negative effect of ethylene on grain size has been confirmed by the application of ethephon (ETH), an ethylene-releasing agent, which decreases grain yield and TGW in wheat, rice, and maize (Ramburan and Greenfield, 2007; Yang *et al*., 2005; Zhou *et al*., 1999). On the other hand, application of ethylene biosynthesis and action inhibitors have the opposite effect. Rice and wheat plants treated with ethylene biosynthesis inhibitors aminoethoxyvinylglycine (AVG), CoCl2 and Co(NO3)2 had higher TGW, grain size, yield and grain filling rate (Beltrano *et al*., 1999; Sarlach *et al*., 2013; Wang *et al*., 2012; Yang J *et al*., 2005). Similarly, foliar spray of ethylene action inhibitors, AgNo3 and 3-cyclopropyl-1-enyl-propanoic acid sodium salt (CPAS) under normal, as well as heat and drought stress conditions significantly enhanced TGW, grain yield and grain filling period of wheat (Huberman *et al*., 2014; Labrana and Araus, 1991). The effect of CPAS on TGW was higher under extreme water stress (Huberman *et al*., 2014) as ethylene production is enhanced by abiotic stress. In their review, Schaller and Binder (2017) noted that the ethylene action inhibitor, 1-Methylcyclopropene (1-MCP) can reduce ethylene responses in various cereal crops with application at booting and/or early flowering stages effective in increasing grain size, TGW, grain filling rate, grain yield, and endosperm cell size under normal (non-stressed) growing conditions. Panda *et al*. (2016) reported that rice treated with 1-MCP under normal field conditions showed a significant improvement in endosperm cell size and area, which is crucial in determining the final grain size. In other studies, 1-MCP treatment merely mitigated the adverse effect of stress on grain size by inhibiting ethylene action. In a heat susceptible cultivar of wheat and in maize, application of 1-MCP during heat stress treatment alleviated the effect of heating on TGW and grain yield (Hays *et al*., 2007). However, the application of 1-MCP was effective in enhancing TGW, grain yield, and grain filling rate in rice with and without stress (Hussain *et al*., 2019; Hussain Sajid *et al*., 2018; Zhang *et al*., 2015).

Due to its gaseous nature resulting in low residues, low toxicity, and existing usage on food crops (Blankenship and Dole, 2003; Muche, 2016) as well as the reported positive effects on grain size, 1-MCP may be a promising plant growth regulator for trialling on teff to improve grain size. However, although a wide variety of evidence indicates that 1-MCP can enhance grain size and yield-related traits in cereals, there is little information on optimal timing of application or the most effective concentration. The major objective of this study was to investigate whether 1-MCP can increase grain size in teff under well-watered or moderate water stress conditions. Secondly, we investigated the optimum application time and concentration of 1-MCP. This is the first study investigating the use of a plant growth regulator to improve grain size in teff. The study therefore also provides important information on the logistics of using plant growth regulators on this semi-domesticated crop with a long flowering period. Finally, the results of the study will also provide useful insight into the role of endogenous ethylene in grain-fill of teff

## Materials and Methods

### Plant material and growth conditions

*Eragrostis tef* seed was obtained from Australian Grain & Forage Seeds (Smeaton, VIC, Australia). The pot experiments were conducted in the University of New England glasshouse during the winter and spring of 2019 and 2020 under natural light and a temperature regime of 23/15°C day/night. Two experiments were conducted: Experiment 1 (2019) involved the treatment of whole plants with 1-MCP; Experiment 2 (2020) involved the treatment of individual panicles. In expt. 1, pots (top and basal diameters 20 and 16 cm respectively, and 19 cm depth) were filled with 3.5 kg of black Dermosol soil (Isbell, 2002) which was collected from the field (−30.4851S, 151.6495E) and fertilised with 47 mg N/kg of dry soil as [NH_4_]_2_ SO_4_, 50 mg K/kg of dry soil as KNO_3_, and 7 mg P/kg of dry soil as [NH_4_]_2_HPO_4_. In expt. 2, pots (top and basal diameters 17 and 14 cm respectively, and 17 cm depth) were filled with 2.6 kg of the same type of soil as expt. 1 and fertilized with 44.9 mg K/kg of dry soil as K_2_SO_4_. No additional N or P was used in expt. 2 as soil analyses indicated the presence of sufficient nutrients.

Pots were watered daily until the seedlings were established (2 to 3 leaves on the main tiller), then at three-day intervals. At the 4 to 6 leaf stage, the plant population was thinned to three plants per pot for expt.1 and one plant per pot for expt.2. In both experiments, half of the plants were maintained at optimal water potential and half were maintained under moderate water stress, applied one week before the expected flowering date based on the data from a preliminary experiment. Water was applied by weight to maintain an optimum matric water potential of −30 kPa, gravimetric soil-water content of 49% as determined from the soil water retention curve (supplementary figure 1). For moderate water stress, water was withheld until the soil reached the target matric water potential of −300 kPa, and gravimetric soil-water content of 37% by measuring the weight of the pot. This water potential for the remainder of the experiment.

### 1-Methylcyclopropene application and experimental procedure

In teff, pollination, panicle emergence, and flowering almost coincide, thus flowering is usually defined as the emergence of the inflorescence tip from the flag leaf sheath (Mengesha and Guard, 1966). Plants were monitored daily during the approach to flowering, with panicles tagged at emergence and the dates of both panicle emergence and completion of panicle flowering recorded (Table 1). The time required for a single panicle to fully emerge ranged from 7 to 15 days, while over the whole plant, panicles emerged over an extended period of 13 to 23 days.

**Table 1.** Recorded phenology data from *Eragrostis tef* plants during each experiment.

| Parameters | Well-watered |  | Moderate water stress |  |
| --- | --- | --- | --- | --- |
|  | Number of days (range) | Number of days (average) | Number of days (range) | Number of days (average) |
| **Experiment One** |  |  |  |  |
| Full emergence of a single panicle | 7 - 10 | 8 | 9 - 12 | 10 |
| Panicle emergence over a whole plant | 15 - 22 | 18 | 15 - 22 | 18 |
| Days to emergence of first and last panicles | 29 - 65 | 50 | 29 - 63 | 48 |
| Days to 50% panicle emergence | 41 - 58 | 47 | 40 - 56 | 46 |
| **Experiment Two** |  |  |  |  |
| Full emergence of a single panicle | 7 - 15 | 10 | 7 - 16 | 11 |
| Panicle emergence over a whole plant | 13 - 23 | 17 | 12 - 22 | 17 |
| Days to emergence of first and last panicles | 33 - 70 | 53 | 33 - 71 | 55 |
| Days to 50% panicle emergence | 48 - 65 | 54 | 36 - 63 | 57 |

The normally gaseous 1-MCP was obtained from Australian Agrofresh (73 Brown Road, VIC, 3809 Australia) as a powdered formulation, SmartFresh™, which has a concentration of 33 g/kg active constituent bound to granules of alpha-cyclodextrin. In expt. 1, 1-MCP was applied to whole plants in plant growth cabinets. The required amount of SmartFresh™ powder to produce a gaseous concentration of 0.1 or 1 PPM in the growth cabinet (Table 2) was added to 5 mL of distilled water. The aqueous solution was sprayed immediately over the plants using an atomizer for 5 seconds before quickly closing the door. Control plants were treated with an equal volume of distilled water at the same time using the same procedures. The air circulation valve of the growth cabinet was closed for 18 hrs following the treatment. Application of 1-MCP was started when at least half of the panicles in each pot had reached flowering. Plants were left in the growth cabinets for a further seven days, with the treatments repeated on the 3^rd^ and 6^th^ days following the first treatment. Plants were then removed back to the glasshouse. As panicle emergence for different tillers of the same plant took place over an extended period, panicles used for data collection were classified in terms of emergence relative to the first application of 1-MCP. Panicles emerging before the application of 1-MCP were designated as PEB, whereas panicles emerging after application of 1-MCP were designated as PEA. Only panicles emerging from 5 days before treatment until 5 days after treatment were used for data collection.

**Table 2.** Treatment Combinations (three 1-MCP concentrations; 0, 0.1, 1 PPM 1-MCP and two moisture treatments; well-watered and moderate water stress) and the required weight of the SmartFresh™ powder to produce the target concentration of 1-MCP for the two successive experiments.

| 1-MCP Concentration (PPM) | Weight (mg) of SmartFresh™ powder for 878 L volume of plant growth-cabinet | Weight (mg) of SmartFresh™ powder for 2 L plastic bag | Moisture Level |
| --- | --- | --- | --- |
| 0 | 0.00 | 0.000 | WW<br>WS |
| 0.1 | 6.39 | 0.015 | WW<br>WS |
| 1 | 63.72 | 0.145 | WW<br>WS |

**Abbreviation;** 1-MCP = 1-Methylcyclopropene, PPM = parts per million, WW = Well-watered, WS = Moderate water stress

In expt. 2, 1-MCP was applied to the first four individual panicles per pot/plant, with the first treatment starting 1 day after panicle emergence. The required amount of SmartFresh™ powder to reach a gaseous concentration of 0.1 or 1 PPM (Table 2) was added to 10 mL of warm distilled water (40-50 ^o^C). 50 μL of this aqueous solution was directly injected into 2 L plastic bags covering individual panicles. Each plastic bag was pre-inflated before injection of the treatment solution using an air compressor and plastic ties. The plastic bags were removed from treated panicles after 24 hrs. Control plants were treated with an equal volume of warm distilled water at the same time using the same procedures. 1-MCP treatments were repeated 3 and 6 days after the initial treatment.

### Data collection

Grains from panicles that emerged from each pot on each day (expt.1) and grains from each panicle (expt.2) were harvested at physiological maturity, threshed, and collected in a sampling bag. The bag was labelled with the date of emergence, pot number, and the number of panicles (only in expt. 1). Whole seeds were oven-dried at 60°C for 24 hrs. A 0.1 g (expt.1) and 0.05 g (expt.2) representative sub-sample of grain from each bag was scanned using an Epson Perfection V700 Photo flatbed scanner (Seiko Epson Corporation, Suwa, Japan) at 1200 dpi to measure grain size parameters. The grain size (area) was determined using image analysis following the method and programs (*SmartGrain*) of Tanabata *et al*. (2012). The program calculates Seed Area (AS), Perimeter Length (PL), Length (L), Width (W), and Length-to-Width ratio (LWR).

Thousand-grain weight was calculated from the seed number (SN) for the 0.1 g or 0.05 g sub-sample. In the treatment of whole plants, yield per panicle was determined by dividing the total grain weight of the panicles that emerged on the same date by the total number of panicles. Following the treatment of individual panicles (expt 2), yield per panicle was determined by weighing the grain mass recovered from each panicle.

The grain yield per plant and biomass yield per plant was determined after harvesting. The harvest index was the ratio of grain yield per plant to biomass yield per plant. Plant height was measured from the base of the plant to the tip of the panicle. Panicle length was measured from the node where the panicle branch starts to the tip of the panicle. Tiller number was determined by the number of tillers per pot/plant.

Time to flowering for each plant was counted as the days from sowing until the emergence of 50% of panicles (Table 1). Days to maturity refers to the time from sowing until physiological maturity and senescence. The grain filling period for each plant was determined as the time from 50% panicle emergence until maturity.

### Experimental design and analysis

Both experiments used a factorial design with three 1-MCP concentrations and two moisture treatments as shown in Table 2. A complete randomized design and a random complete block design were used in expt. 1 and 2, respectively with six replicates per treatment, a replicate consisting of one pot of three plants (expt.1) or one plant (expt.2).

The R-statistical analysis package (R Core Team, 2019) was used to fit generalised linear models and analyses of variance to grain size, TGW, and yield per panicle. 1-MCP application rate, moisture treatment and the date of panicle emergence were used as predictor variables for the analysis of whole plant data. When assessing phenology and yield-related traits, only 1-MCP application rate and moisture treatments were used as predictors. All data from the treatment of individual panicles were analysed with 1-MCP application rate and moisture treatment as predictor variables. Means were compared using post-hoc multiple comparisons (Tukey’s HSD) for the treatment of the whole plant, while the least significant difference test (LSD) was used for the treatment of individual panicle data. In all cases, diagnostic plots were assessed to ensure model assumptions were upheld. Significance was assigned where p≤0.05.

## Results

### Effect of 1-MCP applied to whole plants, on grain size, thousand-grain weight, and yield per panicle

In the first experiment, the effectiveness of 1-MPC application to whole plants was assessed. Thus, panicles on the treated plants were at a variety of ages. The grain size measurements and TGW were assessed on the same sample and the results are very similar; the application of 1-MCP had no overall effect on grain size or TGW across all treatments. However, significant interactions with moisture level and time of application (p = 1.1 × 10^-4^ and 7 × 10^-3^ respectively for grain size, and p = 2 × 10^-4^ and 1 × 10^-3^ for TGW), as well as a significant three-way interaction (p = 0.03 grain size and p = 2 × 10^-3^ TGW), demonstrated that both moisture level and timing of application relative to panicle emergence, altered the response of grains to 1-MCP. The only detectable effect of 1-MCP on grain size was for panicles emerging after the application of 0.1 PPM (Figure 1A). This resulted in an apparent 30% increase in grain size compared to the control plants, when applied to moderately water-stressed plants. However, the application of 1-MCP to well-watered plants did not affect grain size. In contrast, the higher concentration of 1-MCP (1 PPM) increased TGW by 12% compared to the untreated control plant, when applied to well-watered plants and did not have an impact on the TGW of moderately water-stressed plants (Figure 1B). Under both moisture levels, 1-MCP treatment did not influence the grain size or TGW of panicles emerging before the chemical treatment.

**Figure 1.**
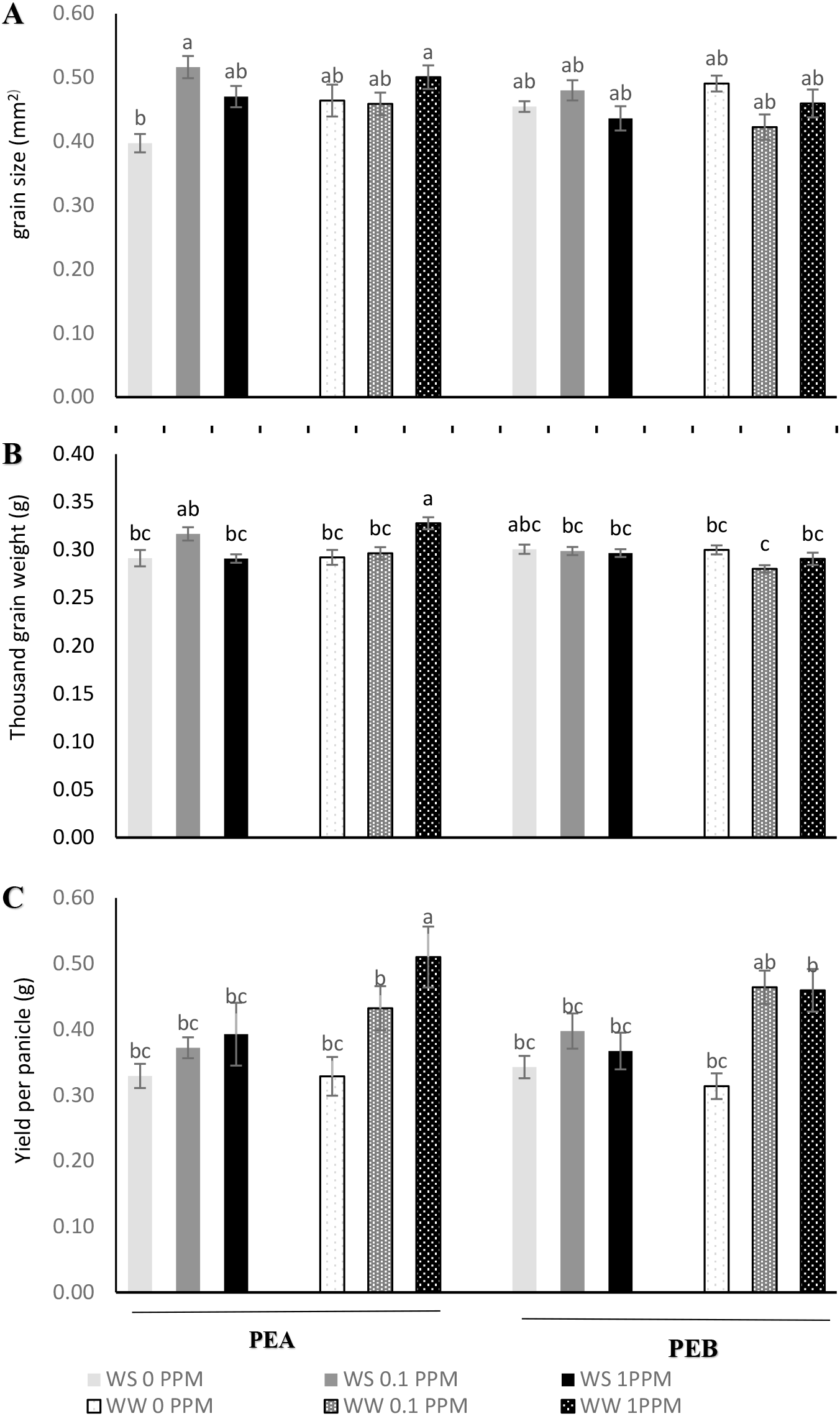
Effect of 1-MCP on Grain size (A), Thousand-grain weight (B), and Yield per panicle (C) of teff with and without water deficit (expt. 1). Data were obtained from panicles emerging between five days before, to five days after the first 1-MCP treatment. Values followed by different letters are significantly different (p ≤ 0.05). Vertical bars represent the mean of 6 replicates ± standard error of the mean. Treatments were; 0 PPM 1-MCP, 0.1 PPM 1-MCP, and 1 PPM 1-MCP; WW = Well-watered and WS = Moderate water stress (applied from one week before flowering until maturity), and PEA = panicle emerged up to 5 days after 1-MCP application and PEB = panicle emerged up to 5 days before 1-MCP application.

Unlike grain size and TGW, the application of 1-MCP did have an overall positive effect on yield per panicle (p = 1.4 × 10^-8^). As above, the interactions of 1-MCP application rate with moisture level (p = 0.002) and time of application (p = 0.01), and three-way interaction (p = 0.04) were all significant, confirming that moisture level and time of 1-MCP application affected the response of the panicles to 1-MCP. At the individual treatment level, the only detectable effect was 1 PPM 1-MCP on yield of well-watered panicles emerging after the treatment (Figure 1C). These plants had a 55% higher yield per panicle than the untreated control plants. However, the application of 1-MCP under moderate water stress did not influence yield per panicle. As was the case for grain size and TGW, the yield of panicles emerging before the treatment was not affected by the application of 1-MCP.

### Effect of 1-MCP applied to whole plants, on grain yield per plant and related traits Grain yield per plant, Biomass yield per plant, and Harvest index

Although there was no detectable effect on grain size, foliar application of 1-MCP had an overall positive effect on grain yield per plant (p = 0.02) and biomass yield per plant (p = 0.03). 1 PPM 1-MCP was the only effective concentration. The interactions of 1-MCP application rate with moisture level for grain yield per plant (p = 0.03) and biomass yield per plant (p = 0.04) were also significant. The effect of 1-MCP was limited to well-watered conditions for both grain and biomass yield per plant (Figure 2A). The well-watered plants treated with 1 PPM 1-MCP had 26% greater grain and biomass yield when compared to the control untreated plant. However, both grain and biomass yield were not affected by the application of 1-MCP under moderate water stress. The harvest index was not affected by the application of 1-MCP (Figure 2B). However, moderately water-stressed plants recorded a higher harvest index (p = 0.002) compared to the control untreated plants.

**Figure 2.**
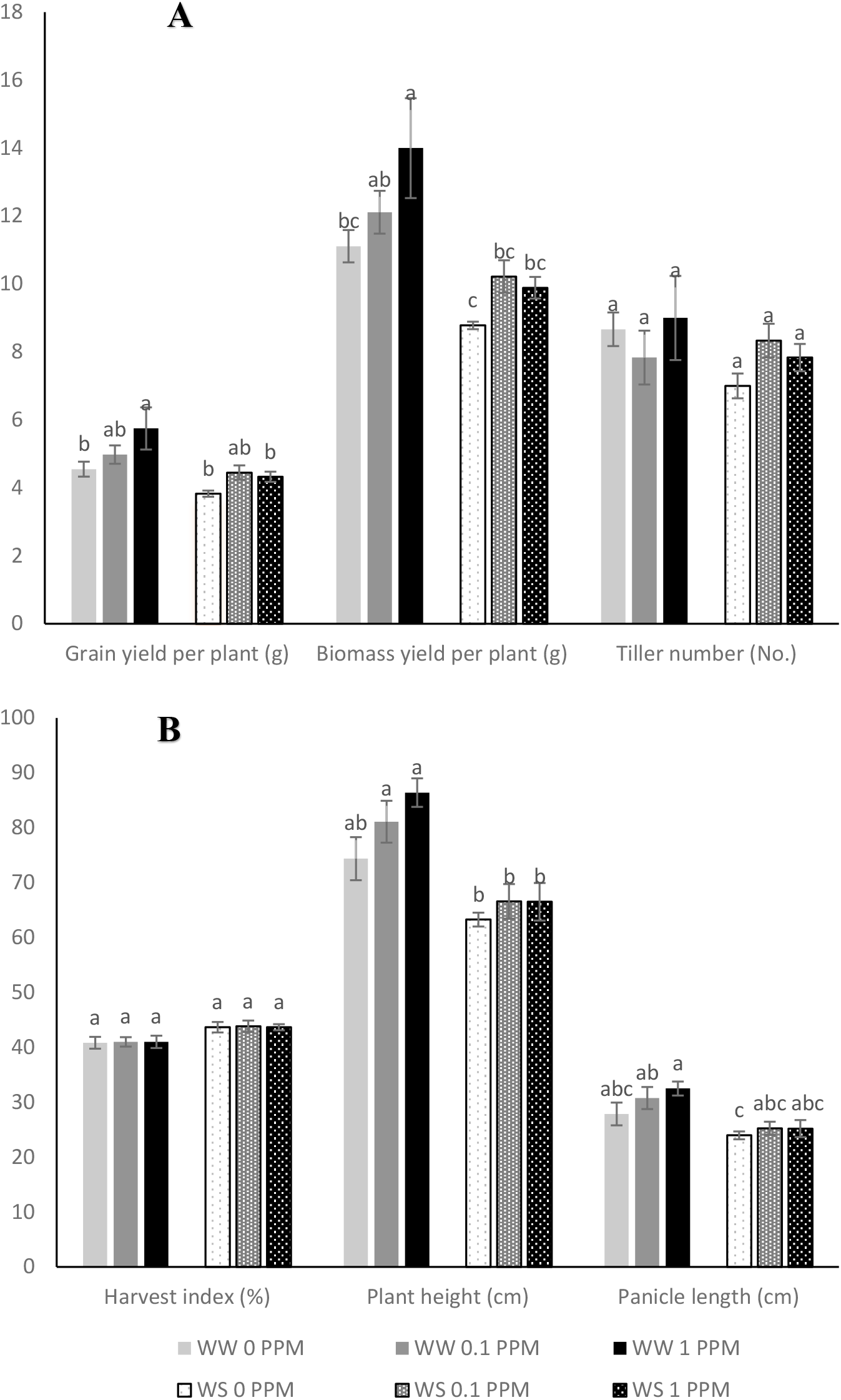
Effects of 1-MCP applied to the whole plant on Grain yield per plant, Biomass yield (g) per plant and Tiller number (**A**), and Harvest index, Plant height and Panicle length (**B**) of teff with and without water deficit (expt. 1). Values followed by different letters are significantly different (p ≤ 0.05). Values are denoted as the mean of six replicates ± standard error of the mean. Where, treatments are; 0 PPM 1-MCP, 0.1 of PPM 1-MCP, and 1 PPM 1-MCP, while WW = Well-watered and WS = Moderate Water stress.

### Plant height, Panicle length, and Tiller number

Application of 1-MCP did not affect plant height and panicle length. However, the interaction between 1-MCP application rate and moisture level was significant for both plant height (p = 0.01) and panicle length (p = 0.03), suggesting moisture level affected the plant reaction to the chemical. Moisture level significantly affected plant height (p = 1.1 × 10^-6^) and panicle length (p = 1.1 × 10^-5^). Well-watered treated plants were taller and had longer panicles than moderately water-stressed treated plants (Figure 2B). However, the tiller number was not significantly affected by 1-MCP application and there was no interaction between the 1-MCP application rate and moisture level.

### Effect of 1-MCP applied to individual panicles, on grain size, thousand-grain weight, and yield per panicle

Although the initial experiment showed no effect of 1-MCP on the main parameters of interest, grain size and TGW, the significant interaction of the treatment with its timing suggested that the variability in panicle age could have been masking a real effect of the treatment. Therefore, the experiment was repeated with the treatment applied to individual panicles of identical developmental age. This time the application of 1-MCP had a significant impact on grain size (p = 1.3 × 10^-12^) and TGW (p = 1.1 × 10^-11^). Treatment with the higher concentration of 1-MCP (1 PPM) resulted in bigger and heavier grains under both moisture levels (Figure 3). There was also a significant interaction between 1-MCP application rate and moisture level (p = 3.5 × 10^-5^ grain size, p = 0.01 TGW), indicating the effect of 1-MCP varied with the moisture level. The effect of 1-MCP was greater under moderate water stress than under well-watered conditions. Compared to the untreated control panicles, the application of 1 PPM 1-MCP increased the grain size of moderately water-stressed panicles by 22%, whereas the grain size of well-watered panicles only increased by 10%. The same treatment also increased TGW by 29% and 16% with and without water deficit, respectively. Grains from moderately water-stressed plants treated with 1-MCP showed an increase in grain size and weight so they were similar to grains from well-watered plants.

**Figure 3.**
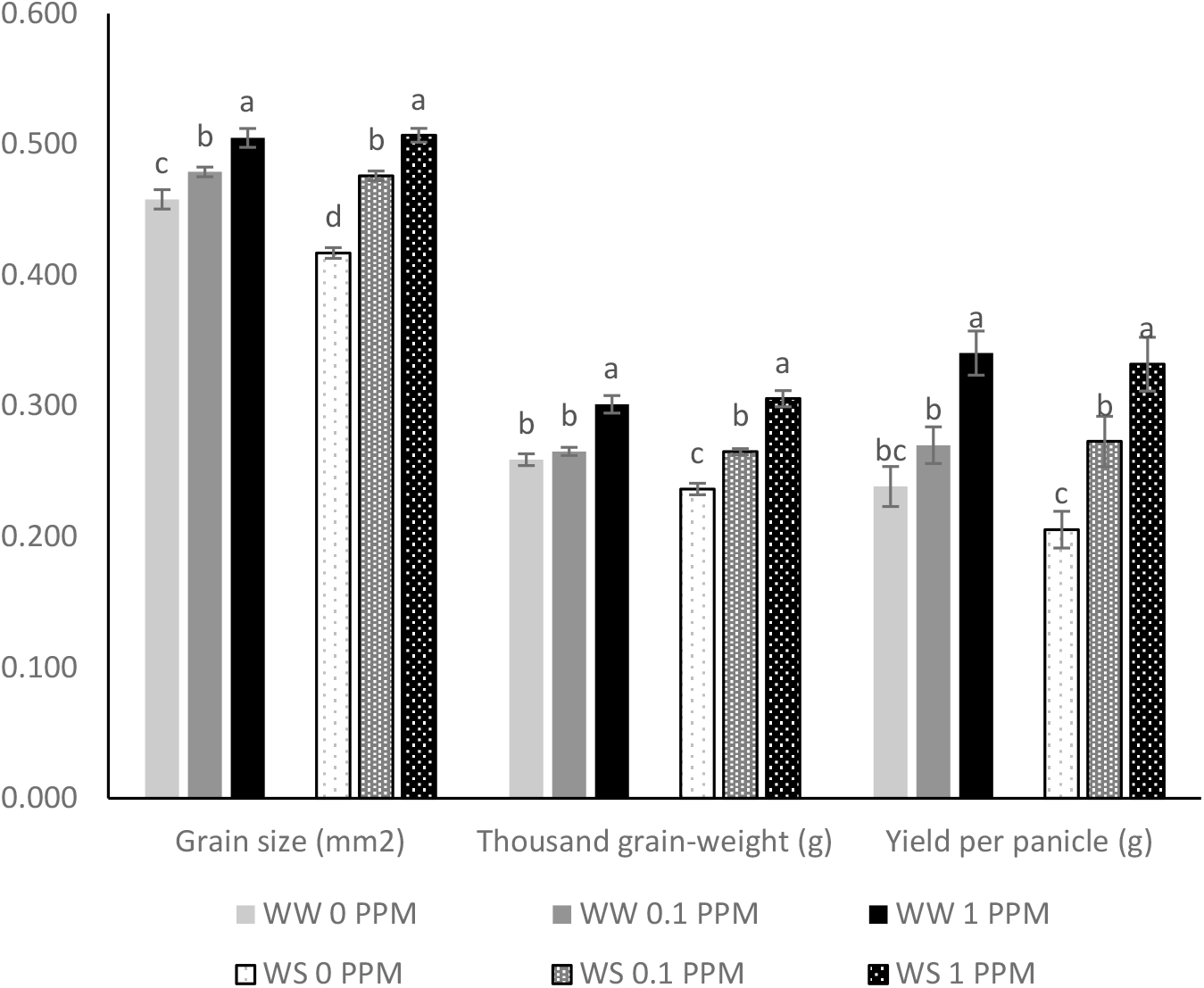
Effect of 1-MCP applied to individual panicles on Grain size, Thousand-grain weight, and Yield per panicle of teff with and without water deficit (expt. 2). Values followed by different letters are significantly different (p ≤ 0.05). Values are denoted as the mean of six replicates ± standard error of the mean. Treatments were; 0 PPM 1-MCP, 0.1 PPM 1-MCP, and 1 PPM 1-MCP; WW = Well-watered and WS = Moderate Water stress (applied from one week before flowering until maturity).

Yield per panicle was determined from the same panicles used to determine grain size and TGW, thus the result is similar. The application of 1-MCP had a positive effect on yield per panicle (p = 7.4 × 10^-8^). As with 1-MCP treatment of whole plants, the higher concentration of 1-MCP (1 PPM) was the most effective, producing the highest yield per panicle (Figure 3). Application of 1 PPM 1-MCP to individual panicles enhanced yield per panicle by 24% and 52% compared with the lower concentration of 1-MCP and untreated control panicles, respectively. However, for this parameter, the interaction between 1-MCP application rate and moisture level was not significant.

### Effect of 1-MCP applied to individual panicles, on grain yield per plant and related traits

Unlike the treatment of the whole plant, the application of 1-MCP to individual panicles had no overall effect on grain yield per plant, biomass yield per plant, tiller number, plant height, panicle length, and harvest index (Figure 4A and B). Each plant/pot had an average of twenty-eight panicles, thus treatment of only four panicles had no significant effect on the whole plant. Similarly, the interaction between 1-MCP application rate and moisture level was not significant for these traits.

**Figure 4.**
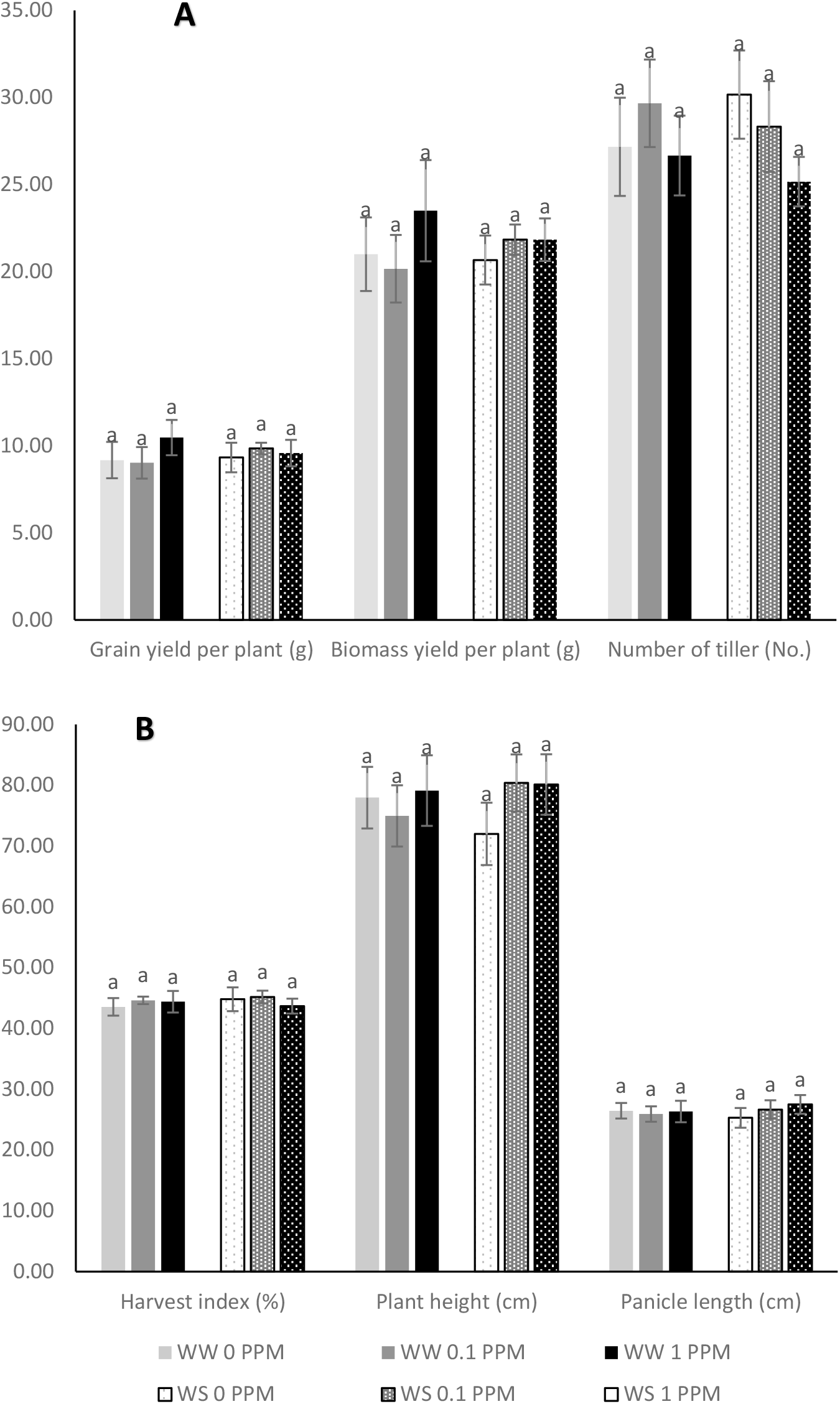
Effects of 1-MCP applied to the first four individual panicles on Grain yield per plant, Biomass yield per plant and Tiller number (**A**), and Harvest index, Plant height and Panicle length (**B**) of teff with and without water deficit (expt. 2). Values followed by different letters are significantly different (p ≤ 0.05). Values are denoted as the mean of six replicates ± standard error of the mean. Treatments were; 0 PPM 1-MCP, 0.1 PPM 1-MCP, and 1 PPM 1-MCP; WW = Well-watered and WS = Moderate Water stress.

### Effect of 1-MCP applied to whole plants and individual panicles on phenology traits

Unlike the other traits described above, 1-MCP applied to the whole plant and/or individual panicle did not have an overall (p ≥ 0.05) effect on phenology traits of teff; days to flowering, days to maturity, and grain filling period (Table 3). The interaction between 1-MCP application rate and moisture level was also not significant.

**Table 3.**
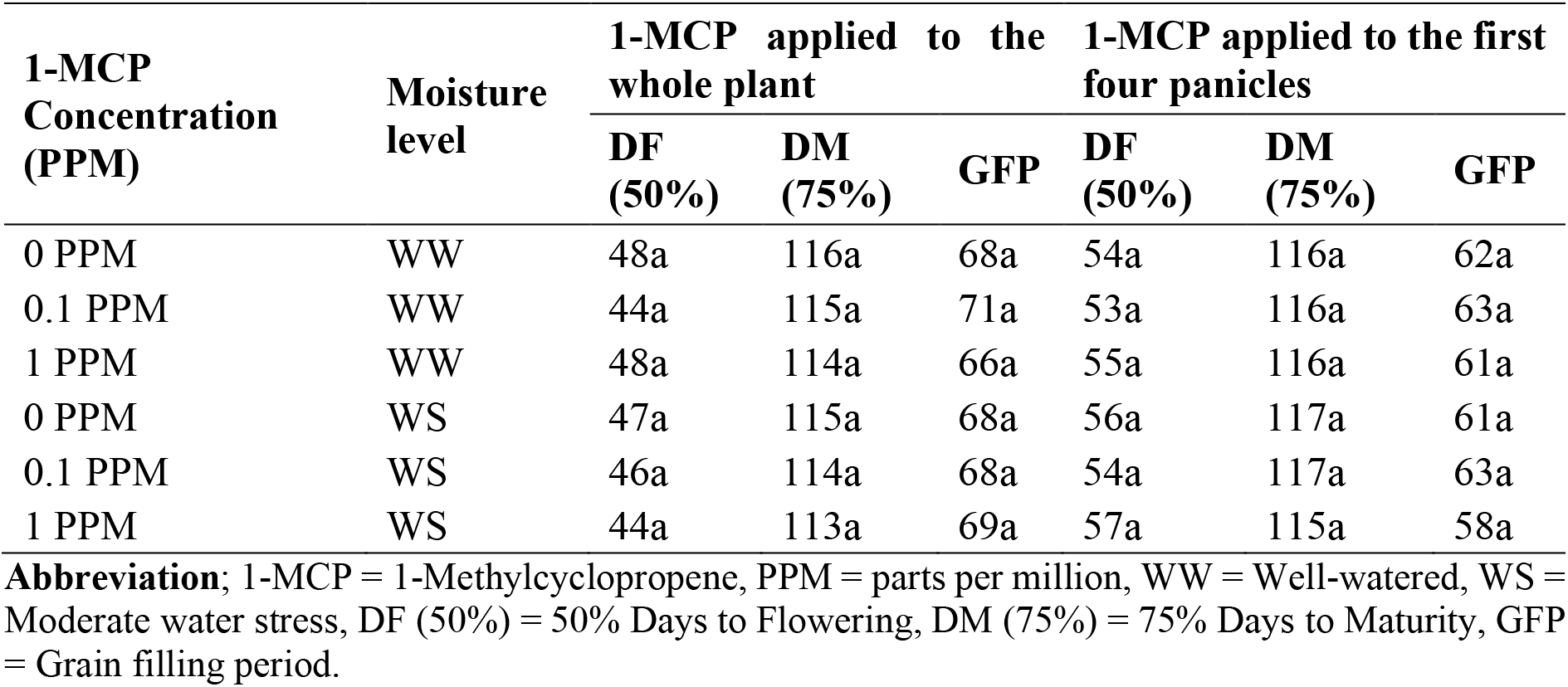
Effect of 1-MCP applied to the whole plant and individual panicles, with and without water deficit on DF (50%), DM (75%), and GFP.

## Discussion

Application of 1-MCP has been used experimentally to improve grain filling rate, TGW, and grain yield in both field and pot experiments on rice, wheat, and maize (Cicchino *et al*., 2013; Hays *et al*., 2007; Hussain Sajid *et al*., 2018; Panda *et al*., 2016). We, therefore, set out to investigate whether treatment with 1-MCP could be used to improve grain size in teff. Our results from the treatment of individual panicles showed that this was indeed the case. In unstressed plants, the grain size increased by 10%, and the TGW by 16%. In plants, grown under moderate water stress (soil matric water potential of −300 KPa), the effect was considerably larger; grain size was increased by 22% and TGW by 29%. The larger effect under water stress was expected as stress has previously been shown to increase ethylene production during early grain fill of other cereals such as wheat, rice, and maize (Cicchino *et al*., 2013; Hays *et al*., 2007; Hussain Sajid *et al*., 2018; Yang W *et al*., 2014). Furthermore, 1-MCP treatment of salt, heat, or water-stressed rice, maize, and wheat during or before grain fill also had a larger positive effect on grain size (Cicchino *et al*., 2013; Hays *et al*., 2007; Hussain *et al*., 2019; Yang *et al*., 2006). Although teff has an excellent capacity to grow and yield in drought-prone areas compared to other crops such as millet, maize, wheat, barley, sorghum, and faba bean (Stallknecht *et al*., 1993; Yadeta *et al*., 2000), it is very sensitive to water stress during the mid-season stage (active grain development) (Yihun *et al*., 2013). When it is exposed to a 25% water deficit the production decreases by about 1 tonne/ha.

Thus, our results demonstrated that 1-MCP treatment has potential to result in meaningful increases in grain size and TGW in teff. On the other hand, our treatment of whole plants did not result in any increase in grain size. The significant interaction between treatment effect and the timing of treatment relative to panicle age indicated that panicles could only respond to the treatment over a narrow window and that the wide range of developmental ages of teff panicles was obscuring any effect on grain size. Detailed studies of ethylene synthesis in developing panicles of rice and wheat have shown that it is maximal at 0 to 4 days post-anthesis, and then gradually decreases (Wang *et al*., 2012; Yang *et al*., 2014). Application of the gaseous ethylene action inhibitor, 1-MCP, is, therefore, most effective within 0 to 4 days of heading. In highly domesticated crops such as rice and wheat, flowering takes place over a narrow window. In wheat, the flowering of individual panicles and the whole plant is completed within a approximately two hours and 3 to 4 days respectively (GRDC, 2018). Similarly in rice, flowering of individual panicles and the whole plant is completed within 5, and 7 to 10 days respectively (Maclean *et al*., 2013). In such crops, 1-MCP can be effective at improving grain size under field conditions. On the other hand, teff is still undergoing the domestication process and has a very different phenology. As shown in table 1, florets on individual panicles emerged and flowered over a 7 to 10 day period, while panicle emergence over a whole plant took 2 to 3 weeks. The three-fold application of 1-MCP to whole plants over 7 days was therefore not effective and field application of 1-MCP treatment is not practical. The phenology of teff is also a contra-indication against possible field use of other PGRs to improve grain size.

In addition, to monitoring grain size and TGW, we also monitored other yield related components including yield per panicle and grain yield per plant as well as harvest index, plant height and panicle length. There was no effect of 1-MCP on harvest index, plant height, tiller number or panicle length. However, there was a substantial increase in yield per panicle of 24% under well-watered conditions and 52% under moderate water stress. As these effects are larger than the effects on grain size and TGW, they indicate that grain number per panicle was also increased. Importantly, there was no decrease in grain number to compensate for the larger grain size. This observation suggests that access to nutrients is not a constraint on grain size.

Although 1-MCP is unlikely to be a practical treatment of field grown teff, the results of this study suggest that genetic manipulation to reduce ethylene production during early grain fill, especially under moderate stress conditions could have a beneficial effect on grain size. However, we suggest that changing the phenology of teff is more likely to be a beneficial strategy for improving yield. Reducing tiller numbers can have multiple benefits in addition to increasing the grain size including reducing lodging and even grain ripening which in turn reduces grain shatter.

## Acknowledgements

We would like to thank Australian Agrofresh for providing 1-MCP and Australian Grain & Forage Seeds for giving teff seed. The authors also acknowledge the UNE for providing a scholarship to Fano Dargo Girmay.

## Author contribution statement

FDG and HMN conceived and designed the research. FDG performed all the experiments and data analysis. RJF helped with the soil laboratory and glasshouse experiment. FDG and HMN wrote, read, and approved the manuscript.

## References

Basunia MA, Nonhebel HM (2019) Hormonal regulation of cereal endosperm development with a focus on rice (Oryza sativa). Functional Plant Biology, 46(6), 493–506. https://doi.org/10.1071/FP18323

Baye K (2014) Teff: nutrient composition and health benefits (Vol. 67): Intl Food Policy Res Inst.

Beltrano J, Ronco MG, Montaldi ER (1999) Drought stress syndrome in wheat is provoked by ethylene evolution imbalance and reversed by rewatering, aminoethoxyvinylglycine, or sodium benzoate. Journal of Plant Growth Regulation, 18(2), 59–64.

Blankenship SM, Dole JM (2003) 1-Methylcyclopropene: a review. Postharvest biology and technology, 28(1), 1–25. https://doi.org/10.1016/S0925-5214(02)00246-6

Chanyalew S, Assefa K, Metaferia G (2013) Phenotypic and molecular diversity in tef. [Achievements and Prospects of Tef Improvement]. 21–31.

Cheng A, Mayes S, Dalle G, Demissew S, et al. (2017) Diversifying crops for food and nutrition security - a case of teff. Biological Reviews, 92(1), 188–198. https://doi.org/10.1111/brv.12225

Cicchino MA, Rattalino Edreira JI, Otegui ME (2013) Maize physiological responses to heat stress and hormonal plant growth regulators related to ethylene metabolism. Crop Science, 53(5), 2135–2146. https://doi.org/10.2135/cropsci2013.03.0136

Crymes AR (2015) The International Footprint of Teff: Resurgence of an Ancient Ethiopian Grain. https://doi.org/10.7936/K70R9MJV

Dereje N, Bekele G, Nigatu Y, Worku Y, et al. (2019) Glycemic Index and Load of Selected Ethiopian Foods: An Experimental Study. Journal of Diabetes Research, 2019. https://doi.org/10.1155/2019/8564879

FAO (2018) Food loss analysis: causes and solutions - Case study on the teff value chain in the Federal Democratic Republic of Ethiopia. Rome. 48 pp. Licence: CC BY-NC-SA 3.0 IGO. Retrieved from http://www.fao.org/3/I9972EN/i9972en.pdf.

Fufa B, Behute B, Benesh K, Simons R, et al. (2013) Analysis of Tef Value Chain in Ethiopia. Achievements and Prospects of Tef Improvement, 305–322.

Gebru YA, Hyun-Ii J, Young-Soo K, Myung-Kon K, et al. (2019) Variations in Amino Acid and Protein Profiles in White versus Brown Teff (Eragrostis Tef) Seeds, and Effect of Extraction Methods on Protein Yields. Foods, 8(6), 202. https://doi.org/10.3390/foods8060202

GRDC (Producer). (2018). Plant growth (phenology). Wheat development stages. Retrieved from https://grdc.com.au/data/assets/pdf_file/0023/366251/GrowNote-Durum-West-3-Plant-Growth.pdf

Hays DB, Do JH, Mason RE, Morgan G, et al. (2007) Heat stress induced ethylene production in developing wheat grains induces kernel abortion and increased maturation in a susceptible cultivar. Plant Science, 172(6), 1113–1123. https://doi.org/10.1016/j.plantsci.2007.03.004

Huberman M, Riov J, Goldschmidt EE, Apelbaum A, et al. (2014) The novel ethylene antagonist, 3-cyclopropyl-1-enyl-propanoic acid sodium salt (CPAS), increases grain yield in wheat by delaying leaf senescence. Plant Growth Regulation, 73(3), 249–255. https://doi.org/10.1007/s10725-013-9885-5

Hussain S, Bai ZG, Huang J, Cao XC, et al. (2019) 1-Methylcyclopropene Modulates Physiological, Biochemical, and Antioxidant Responses of Rice to Different Salt Stress Levels. Frontiers in Plant Science, 10, 18. https://doi.org/10.3389/fpls.2019.00124

Hussain S, Zhong C, Bai Z, Cao X, et al. (2018) Effects of 1-Methylcyclopropene on Rice Growth Characteristics and Superior and Inferior Spikelet Development Under Salt Stress. Journal of Plant Growth Regulation, 1–17. https://doi.org/10.1007/s00344-018-9800-4

Isbell RF (2002) The Australian soil classification: Revised edition, CSIRO Publishing, http://hdl.handle.net/102.100.100/198257?index=1.

Kesavan M, Song JT, Seo HS (2013) Seed size: a priority trait in cereal crops. Physiologia Plantarum, 147(2), 113–120. https://doi.org/10.1111/j.1399-3054.2012.01664.x

Ketema S (1997) Tef-Eragrostis Tef (Zucc.) (Vol. 12): Bioversity International.

Labrana X, Araus JL (1991) Effect of foliar applications of silver-nitrate and ear removal on carbon-dioxide assimilation in wheat flag leaves during grain-filling. Field Crops Research, 28(1-2), 149–162. https://doi.org/10.1016/0378-4290(91)90080-F

Li L, Gu W, Zuo S, Meng Y, et al. (2020) Effects of thidiazuron and ethephon on the grain filling and dehydration characteristics of maize in Northeast China. Archives of Agronomy and Soil Science, 1–17. https://doi.org/10.1080/03650340.2020.1858480

Maclean J, Hardy B, Hettel G (2013) Rice Almanac: Source book for one of the most important economic activities on earth: IRRI.

Mengesha MH, Guard A (1966) Development of the embryo sac and embryo of teff, Eragrostis tef. Canadian Journal of Botany, 44(8), 1071–1075. https://doi.org/10.1139/b66-114

Muche BM (2016) Effect of 1-methylcyclopropene (1-MCP) on the Flavour Metabolites of Apple Juice. (PHD), DALHOUSIE UNIVERSITY, dalspace.library.dal.ca.

Panda B, Badoghar A, Sekhar S, Shaw B, et al. (2016) 1-MCP treatment enhanced expression of genes controlling endosperm cell division and starch biosynthesis for improvement of grain filling in a dense-panicle rice cultivar. Plant Science, 246, 11–25. https://doi.org/10.1016/j.plantsci.2016.02.004.

Panda BB, Sekhar S, Dash SK, Behera L, et al. (2018) Biochemical and molecular characterisation of exogenous cytokinin application on grain filling in rice. Bmc Plant Biology, 18(1), 89. https://doi.org/10.1186/s12870-018-1279-4

R Core Team (2019) R: A Language and Environment for Statistical Computing In. R Foundation for Statistical Computing, Vienna, Austria: https://www.R-project.org/.

Ramburan S, Greenfield P (2007) The effects of chlormequat chloride and ethephon on agronomic and quality characteristics of South African irrigated wheat. South African Journal of Plant and Soil, 24(2), 106–113. https://doi.org/10.1080/02571862.2007.10634790

Sarlach R, Bains N, Gill M (2013) Effect of Foliar Application of Osmoprotectants and Ethylene Inhibitor to Enhance Heat Tolerance in Wheat (Triticum aestivum L.). Vegetos-An International Journal of Plant Research, 26(1), 266–271. https://doi.org/10.5958/j.2229-4473.26.1.038

Schaller GE, Binder BM (2017) Inhibitors of Ethylene Biosynthesis and Signaling. Ethylene Signaling, 223–235. https://DOI:10.1007/978-1-4939-6854-1_15

Sivakumar T, Virendranath, Srivastava GC (2001) Effects of Benzyl Adenine and Abscisic Acid on Grain Yield and Yield Components in Triticale and Wheat. Journal of Agronomy and Crop Science, 186(1), 43–46. https://doi.org/10.1046/j.1439-037x.2001.00450.x

Spaenij-Dekking L, Kooy-Winkelaar Y, Koning F (2005) The Ethiopian cereal tef in celiac disease. New England Journal of Medicine, 353(16), 1748–1749. https://doi.10.1056/NEJMc051492

Stallknecht GF, Gilbertson KM, Eckhoff J (1993). Teff: Food crop for humans and animals. p. 231–234. In: J. Janick and J.E. Simon (eds.), New crops. Wiley, New York. Retrieved from https://hort.purdue.edu/newcrop/proceedings1993/V2-231.html

Tanabata T, Shibaya T, Hori K, Ebana K, et al. (2012) SmartGrain: high-throughput phenotyping software for measuring seed shape through image analysis. Plant physiology, 160(4), 1871–1880. https://doi.org/10.1104/pp.112.205120

Van Delden S, Vos J, Ennos A, Stomph T (2010) Analysing lodging of the panicle bearing cereal teff (Eragrostis tef). New Phytologist, 186(3), 696–707. https://doi.org/10.1111/j.1469-8137.2010.03224.x

Vavilov NI (1951) The origin, variation, immunity and breeding of cultivated plants: Soil Science 72(6):p 482.

Wang F, Cheng F, Zhang G (2006) The relationship between grain filling and hormone content as affected by genotype and source–sink relation. Plant Growth Regulation, 49(1), 1–8. https://doi.org/10.1007/s10725-006-0017-3

Wang ZQ, Xu YJ, Wang JC, Yang JC, et al. (2012) Polyamine and ethylene interactions in grain filling of superior and inferior spikelets of rice. Plant Growth Regulation, 66(3), 215–228. https://doi.org/10.1007/s10725-011-9644-4

Wei S, Wang X, Li G, Qin Y, et al. (2019) Plant density and nitrogen supply affect the grain-filling parameters of maize kernels located in different ear positions. Frontiers in Plant Science, 10, 180. https://doi.org/10.3389/fpls.2019.00180

Yadeta K, Ayele G, Negatu W (2000). Farming Systems Research on Tef: Smallholders“ Production Practices. Paper presented at the 2000). Narrowing the Rift: Tef Research and Development. Proceedings of the International Workshop on Tef Genetics and Improvement, Debrezeit, Ethiopia.

Yang J, Zhang J, Wang Z, Liu K, et al. (2005) Post-anthesis development of inferior and superior spikelets in rice in relation to abscisic acid and ethylene. Journal of Experimental Botany, 57(1), 149–160. https://doi.org/10.1093/jxb/erj018

Yang J, Zhang J, Wang Z, Zhu Q (2003) Hormones in the grains in relation to sink strength and postanthesis development of spikelets in rice. Plant Growth Regulation, 41(3), 185–195. https://doi.org/10.1023/B:GROW.0000007503.95391.38

Yang JC, Zhang JH, Liu K, Wang ZQ, et al. (2006) Abscisic acid and ethylene interact in wheat grains in response to soil drying during grain filling. New Phytologist, 171(2), 293–303. https://doi.org/10.1111/j.1469-8137.2006.01753.x

Yang JC, Zhang JH, Ye YX, Wang ZQ, et al. (2004) Involvement of abscisic acid and ethylene in the responses of rice grains to water stress during filling. Plant Cell and Environment, 27(8), 1055–1064. https://doi.org/10.1111/j.1365-3040.2004.01210.x

Yang W, Yin Y, Li Y, Cai T, et al. (2014) Interactions between polyamines and ethylene during grain filling in wheat grown under water deficit conditions. Plant Growth Regulation, 72(2), 189–201. https://doi.org/10.1007/s10725-013-9851-2

Yihun YM, Haile AM, Schultz B, Erkossa T (2013) Crop water productivity of irrigated teff in a water stressed region. Water resources management, 27(8), 3115–3125. https://doi.org/10.1007/s11269-013-0336-x.

Zhang JH, Lin YJ, Zhu LF, Yu SM, et al. (2015) Effects of 1-methylcyclopropene on function of flag leaf and development of superior and inferior spikelets in rice cultivars differing in panicle types. Field Crops Research, 177, 64–74. https://doi.org/10.1016/j.fcr.2015.03.003

Zhang WY, Cao ZQ, Zhou Q, Chen J, et al. (2016) Grain Filling Characteristics and Their Relations with Endogenous Hormones in Large-and Small-Grain Mutants of Rice. Plos One, 11(10), 20. https://doi.org/10.1371/journal.pone.0165321

Zhou X, Mackenzie A, Madramootoo C, Smith D (1999) Effects of stem-injected plant growth regulators, with or without sucrose, on grain production, biomass and photosynthetic activity of field-grown corn plants. Journal of Agronomy and Crop Science, 183(2), 103–110. https://doi.org/10.1046/j.1439-037x.1999.00331.x

