## Supplementary Figure 1. for "The Effect of 1-MCP on grain size and yield-related traits of *Eragrostis tef*"

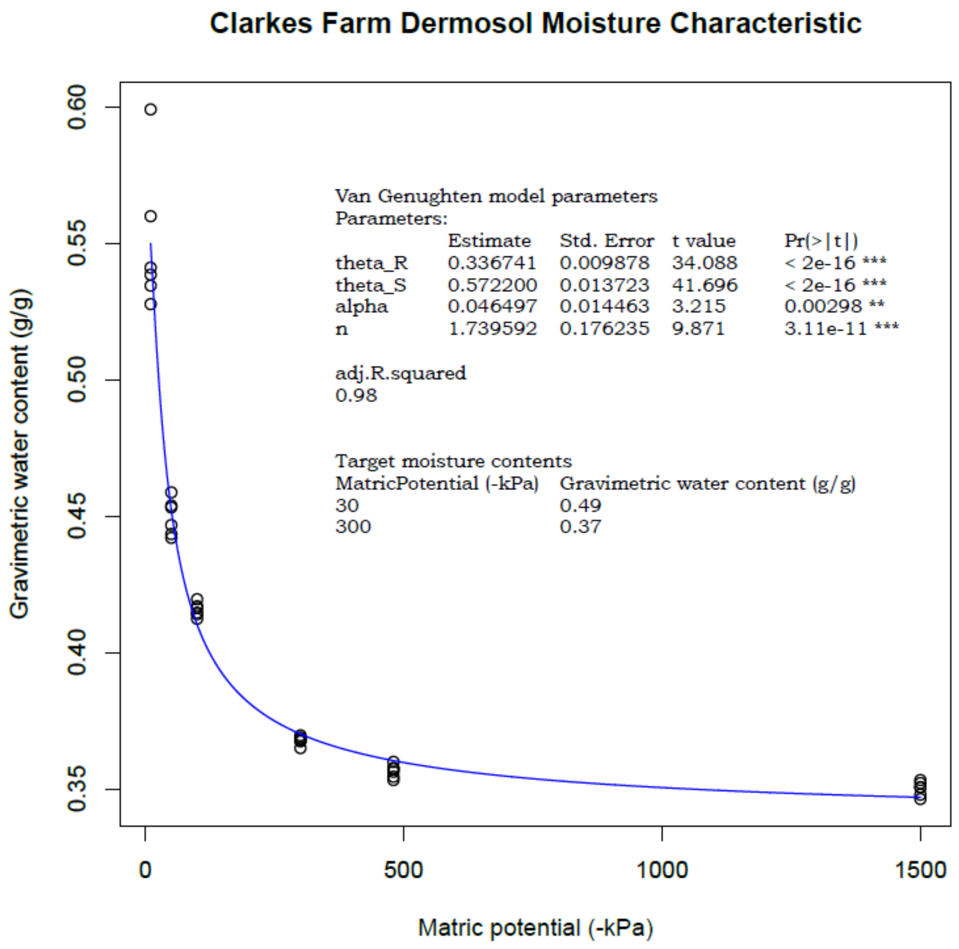


**Supplementary Figure 1.** The relationship between matric potential and gravimetric water content (water retention curve).
